# The host transcriptional response to superinfection by influenza virus and streptococcus pneumonia

**DOI:** 10.1101/2022.08.15.503953

**Authors:** Ofir Cohn, Gal Yankovitz, Michal Mandelboim, Naama Peshes-Yaloz, Eran Bacharach, Irit Gat-Viks

## Abstract

Secondary bacterial challenges during influenza virus infection (‘superinfection’) cause excessive mortality and hospitalization. Here we present a longitudinal study of gene-expression changes in murine lungs during superinfection, with an initial influenza A virus (IAV) infection and a subsequent *Streptococcus pneumonia* (SP) infection. In addition to the well-characterized impairment of the innate immune response, we identified superinfection-specific alterations in endothelial-related genes, including a previously uncharacterized rapid downregulation of particular angiogenic and vascular markers. Superinfection-specific alterations were also evident in the analysis of cellular states related to the host’s immune resistance against pathogens. We found that superinfected mice manifested an excessive rapid induction of immune resistance starting only a few hours after the secondary bacterial challenge. In addition, there was a substantial rewiring of the resistance program: interferon-regulated genes were switched from positive to negative correlations with resistance, whereas genes of fatty-acid metabolism were switched from negative to positive correlations with resistance. Thus, the transcriptional resistance state in superinfection is reprogrammed toward repressed interferon signaling and induced fatty acid metabolism. Our findings suggest new insights into the remodeling of the host defense upon superinfection, providing promising targets for future therapeutic interventions.

## Introduction

Since the dawn of human history, infectious diseases have placed a significant burden on public health and the global economy (Global burden disease study, 2018), with respiratory infections constituting a leading cause of morbidity and mortality worldwide, particularly in immunocompromised subpopulations such as young children and older adults (Kunisaki and Janoff, 2009). Among respiratory viral pathogens, the influenza A virus (IAV) poses a continual threat to global health as it is responsible for seasonal outbreaks and occasional pandemics in humans, contributing to 3–5 million cases of severe disease and 290,000 to 650,000 deaths annually (Johnson and Mueller, 2002), with an increased mortality rate during pandemic years (Chien et al., 2009). This enhanced pathogenesis is associated with secondary bacterial pneumonia, attributing to 40-95% of IAV-related mortality during past pandemics (Morens et al., 2004). *Streptococcus pneumonia* (SP) is a commonly identified bacterium in IAV pandemics and a prominent etiological agent of secondary bacterial pneumonia. As suggested by clinical and autopsy examinations, during the “Spanish” IAV pandemic in 1918–1919, more than 95% of the deaths (~50 million) were complicated by bacterial infections, most commonly by SP (MacIntyre et al., 2018, p. 09; Morens et al., 2008). The combination of viral infection (e.g., of IAV) and secondary bacterial infection is called ‘superinfection’.

Several physical and immunological mechanisms explain the increased permissiveness of IAV-infected lungs to subsequent bacterial infections. Particularly, various studies highlighted virus-induced damage of the respiratory epithelium as a primary factor for increased bacterial adherence and replication, resulting in a loss of lung repair processes and impaired function (Brian W. Donnelly et al., 1990; Kash et al., 2011; Nugent and Pesanti, 1983; Plotkowski et al., 1986). In addition, multiple studies have revealed IAV-infection-dependent reduction in innate immune activity, such as depletion or dysfunction of macrophages and neutrophils that are essential for early bacterial clearance (Ghoneim et al., 2013; McNamee and Harmsen, 2006). This dysfunction, in turn, leads to elevated production of cytokines and chemokines (Shahangian et al., 2009; Sun and Metzger, 2008; van der Sluijs et al., 2004) that subsequently affect other immune cell types (e.g., T-cells, monocytes) (Ellis et al., 2015; Li et al., 2012).

Global gene-expression analyses allow an unbiased genome-scale discovery of candidate mechanisms. However, until now, global gene-expression studies of superinfections have primarily been restricted to analyzing a single pre-selected time point (Kash et al., 2011; Luo et al., 2017), thereby providing a limited ability to pinpoint reproducible and dynamic alterations associated with superinfection. To better understand superinfection-specific changes, here we employed a superinfection model consisting of an SP infection that follows an initial IAV infection. Particularly, we have analyzed longitudinal transcriptional responses during the IAV/SP superinfection compared to IAV-only infection, SP-only infection, and control (PBS) treatments.

## Results

### Superinfection is associated with a severe disease course

To investigate superinfection, we followed a model in which mice are exposed to SP 5 days after the initial IAV infection (*n* = 16)(Schliehe et al., 2015; Shahangian et al., 2009). We compared the IAV/SP superinfection to three control treatments: IAV-only infection (IAV on day 1, PBS on day 5, *n*=18), SP-only infection (PBS on day 1, SP on day 5, *n*=7), and a mock-infection control (PBS on days 1 and 5, *n*=10) (**Figure 1A**). Several lines of evidence demonstrated the enhanced severity of the superinfection compared to the other treatments. First, we found that superinfected mice showed increased gradual weight loss compared to the three control groups (**Figure 1B**). Second, superinfected mice had a higher bacterial load in the lungs than those infected only with bacteria (**Figure 1C**; the bacterial load was measured using 16S rRNA by quantitative RT-PCR (qRT-PCR)). Third, we quantified the viral burden in the lungs by measuring lung transcriptomes that captured both the host and the viral RNA. The viral burden in superinfected mice was similar to or higher than the top viral burden in the IAV-only group (**Figure 1D**). Thus, consistent with previous studies (McCullers and Rehg, 2002; Sharma-Chawla et al., 2016), IAV/SP superinfection has a substantial severity compared to IAV-only and SP-only infections.

**Figure 1.**
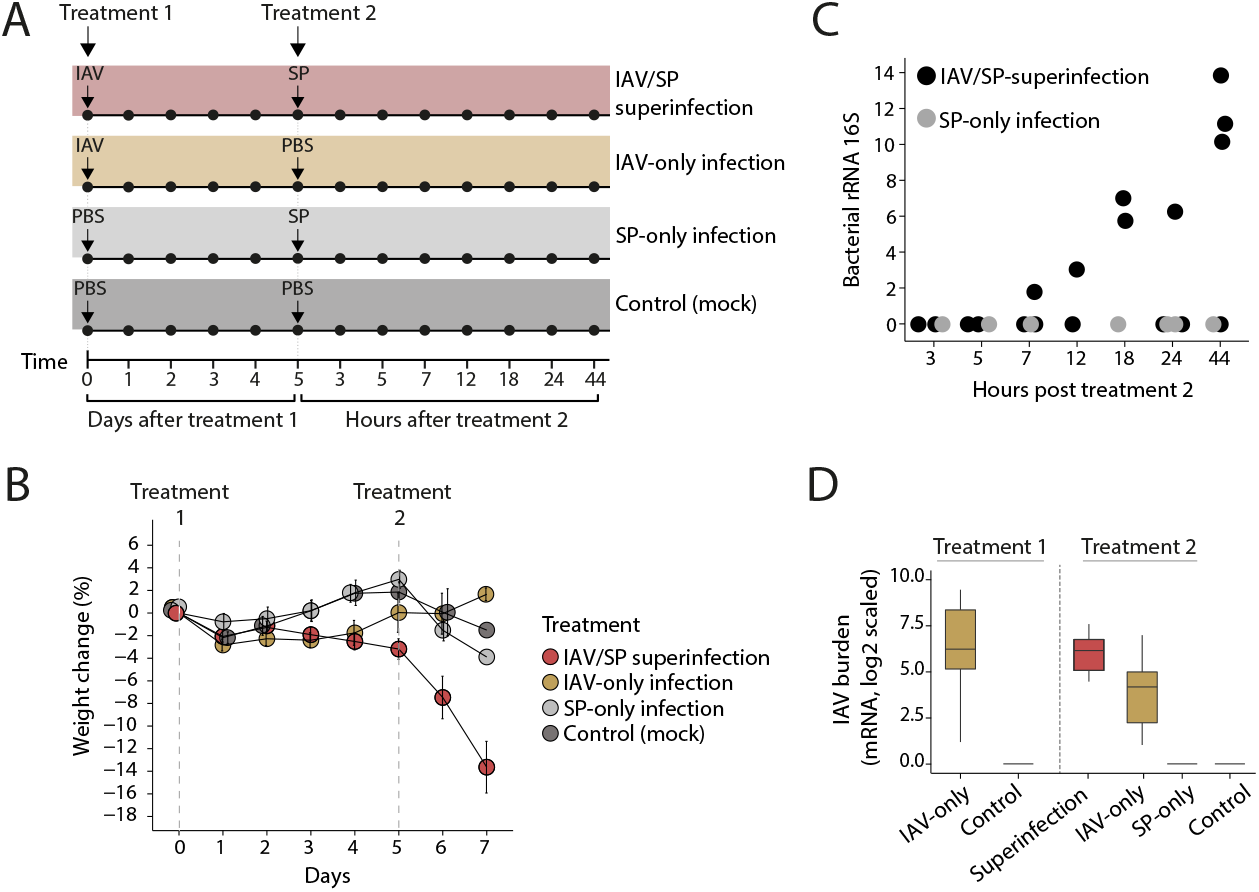
High disease severity in superinfection with IAV and SP. (**A**) Schematics of the study design and timelines for IAV-only infection, SP-only infection, IAV/SP superinfection, and a mock (PBS) control treatment. (**B**) Mean percent weight loss from baseline ± standard error, for each treatment at indicated days. (**C**) Quantification by qPCR of bacterial 16S rRNA gene copies in lungs of superinfected and SP-only infected mice at indicated hours after secondary bacterial infection. (**D**) Quantification of viral burden in lungs, based on viral mRNA levels (log scaled). Presented is the mean viral mRNA level ± standard error for each treatment group, during days 1 to 5 after treatment 1 (left) and 3-44 hours after treatment 2 (right).

### The IAV/SP superinfection is associated with massive induction and repression of genes

Given the limited knowledge about the factors responsible for the increased host volunrability upon superinfection, we aimed to identify host transcriptional changes specifically manifested in the secondary infection. As an initial indication, we observed that the overall transcriptional state is distinct in IAV/SP superinfection compared to the other treatments (**Figure 2A**). Particularly, we observed a clear separation along the first principle component (PC1) axis between the IAV/SP superinfection treatment and the other treatments.

**Figure 2.**
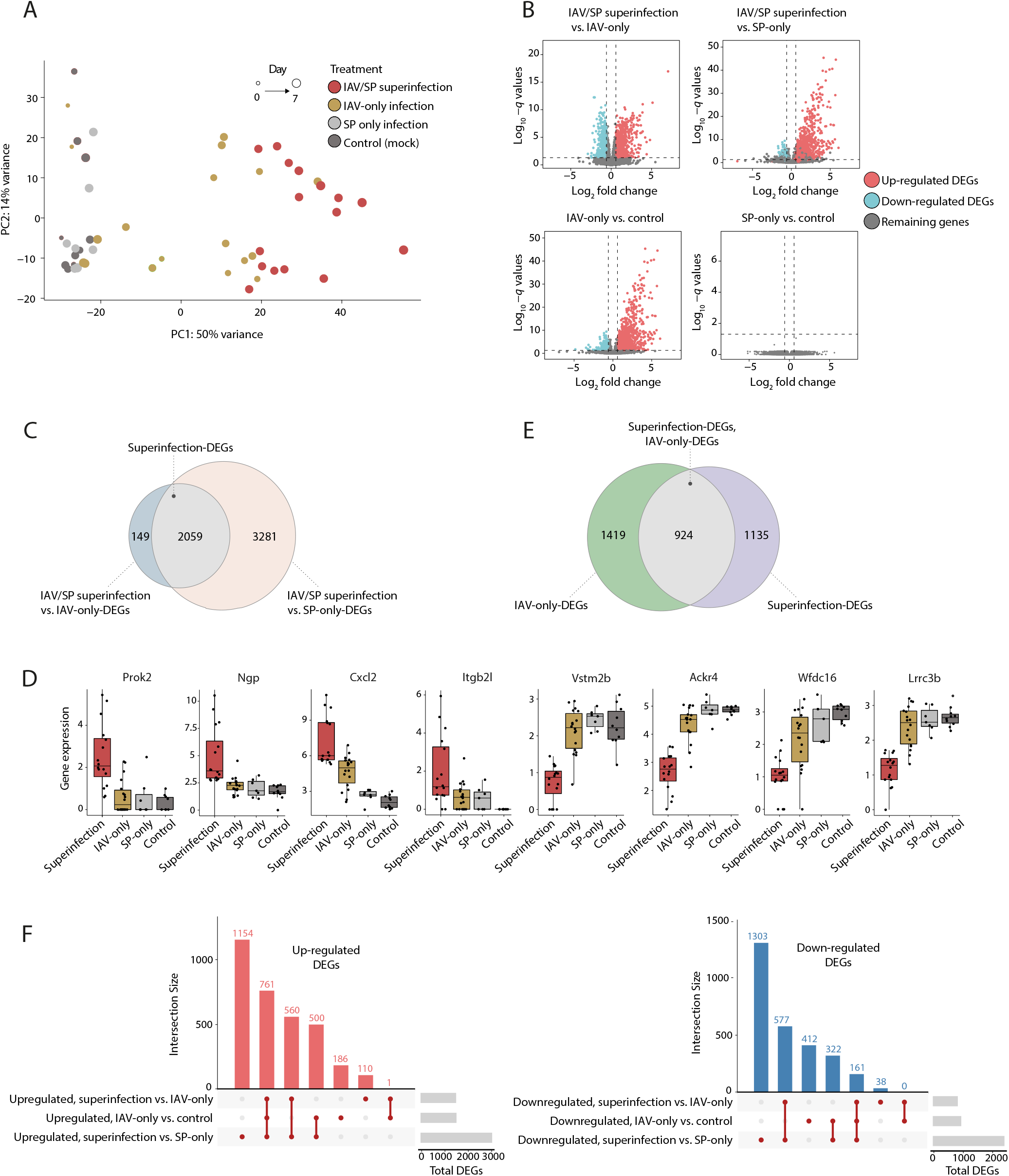
Massive superinfection-specific induction and repression of genes. (**A**) IAV/SP superinfection is associated with a global change in the overall gene expression state. Principal component analysis (PCA) of transcriptome profiles of mice from the four treatment groups. For each mouse (a circle), its treatment group is color coded, and the size of its circle corresponds to the time (days) post the start of experiment. (**B**) Volcano plots indicating differentially expression genes (DEGs) of the compared treatment groups (each dot represents a single gene). DEGs are indicated in blue (down-regulated) or red (up-regulated). The remaining genes are indicated in grey. (**C**) Venn diagram displaying the definition of superinfection-DEGs: genes that are differentially expressed in comparison of IAV/SP superinfection against each of the two single infections. (**D**) Selected superinfection-DEGs. Presented are four top upregulated genes and four top downregulated genes in each of the compared groups. Individual samples are shown as dots. © Venn diagram displaying the intersection between two sets: superinfection-DEGs (defined in C) and IAV-DEGs (IAV-only versus control). (**F**) Upset plot visualizing the intersections of three sets of DEGs: superinfection versus IAV-only, superinfection versus SP-only, and IAV-only versus control. Left/right: each gene set includes only the upregulated/downregulated genes. In all plotsDEGs are genes with *q*-value < 0.05 and FC>1.5.

When we compared IAV/SP superinfection to IAV-only samples, a total of 2208 differentially expressed genes (DEGs) were detected. Of which, 1432 genes (65%) were significantly upregulated and 776 genes (35%) were significantly downregulated (using a *q* value < 0.05 and fold change [FC] > 1.5 cutoffs; **Figure 2B**). Importantly, 93% (2059) of the genes that were differentially expressed in superinfection compared to IAV-only infection were also differentially expressed in superinfection compared to SP-only infection (*q* value < 0.05 and FC > 1.5; hypergeometric [HG] p<<10^-100^; see overlaps in **Figure 2C** and **Supp Table 1**). We refer to these 2059 genes as ‘superinfection-DEGs’, including 1321 upregulated and 738 downregulated genes (exemplified in **Figure 2D**). The massive number of genes primarily respond to superinfection rather than IAV-only or bacteria-only infections suggests a substantial transcriptional response during the secondary infection.

Superinfection-DEGs define a differential expression between superinfection and a single (IAV or SP) infection. Next, we asked whether these superinfection-DEGs are differentially expressed when a single infection is compared to mock infection. In the following, we first describe the SP-only experiment compared to mock and then the IAV-only experiment compared to mock. We found only one DEG for SP-only treatment compared to mock infection (*q* value ≤ 0.05 and FC > 1.5, denoted ‘SP-DEG’), implying that the superinfection response is not observed in the SP-only treatment. In contrast, there was an extensive change in expression following IAV-only treatment compared to mock-infection treatment (2343 DEGs, *q* value ≤ 0.05 and FC > 1.5, denoted ‘IAV-DEGs’; **Figure 2E**). Many of the superinfection-DEGs were also IAV-DEGs (924 of 2059 genes, HG p<<10^-100^). Out of these 924 genes, 922 (>99%) responded in the same directions as superinfection-DEGs and IAV-DEGs. However, there is a clear difference between upregulated and downregulated superinfection-DEGs: among 1321 upregulated superinfection-DEGs, 761 genes (57%) were also upregulated IAV-DEGs, whereas among 738 downregulated superinfection-DEGs, only 161 (21%) were also downregulated in the IAV-only treatment (*p* <10^-5^, χ^2^ test; **Figure 2F**). Thus, the extensive downregulation is a unique property of superinfection; in contrast, part of the upregulation in superinfection is an amplification of the initial transcriptional response to IAV infection.

We next focused on the top superinfection-DEGs, for which the differences between superinfection and single infections are maximized (158 upregulated and 129 downregulated superinfection-DEGs, **Methods**). As shown in **Figure 3A**, the top downregulated and upregulated genes represent a rapid change in the transcripts levels within several hours post-secondary bacterial infection. Functional analysis revealed that the downregulated superinfection-DEGs are involved in angiogenesis and vascular-associated terms, such as ‘regulation of blood vessel endothelial cell proliferation’ (*p* <10^-5^, HG test) and ‘morphogenesis of an endothelium’ (*p* <10^-5^, HG test) (**Figure 3B**; see exemplified genes in **Figure 3C**). In agreement, vascular-related functions from the Ingenuity annotation are mainly enriched in the superinfection-DEGs compared to the enrichment in the IAV-DEGs (**Figure 3D**). The top upregulated superinfection-DEGs are involved in immune responses such as inflammation (HG test *p* <10^-20^) and cytokine production (HG test *p* <10^-12^) – e.g., *Il10, Fcgr3, and Ccr1* (**Figure 3B,C**). Therefore, our findings highlight endothelial and vascular dysregulation, which could be related to the documented increased risk of barrier disruption, pulmonary thrombosis, and excessive vascular leak (Sender Vicky et al., 2020; Yang and Tang, 2016). Of note, whereas some of the top superinfection-DEGs are well established (e.g., *Cxcl2* (Shirey Kari Ann et al., n.d.)), many other superinfection-DEGs are not yet reported, such as *Ngp, Ackr4, Vstm2b* and *Prok2* (**Figure 2D**).

**Figure 3.**
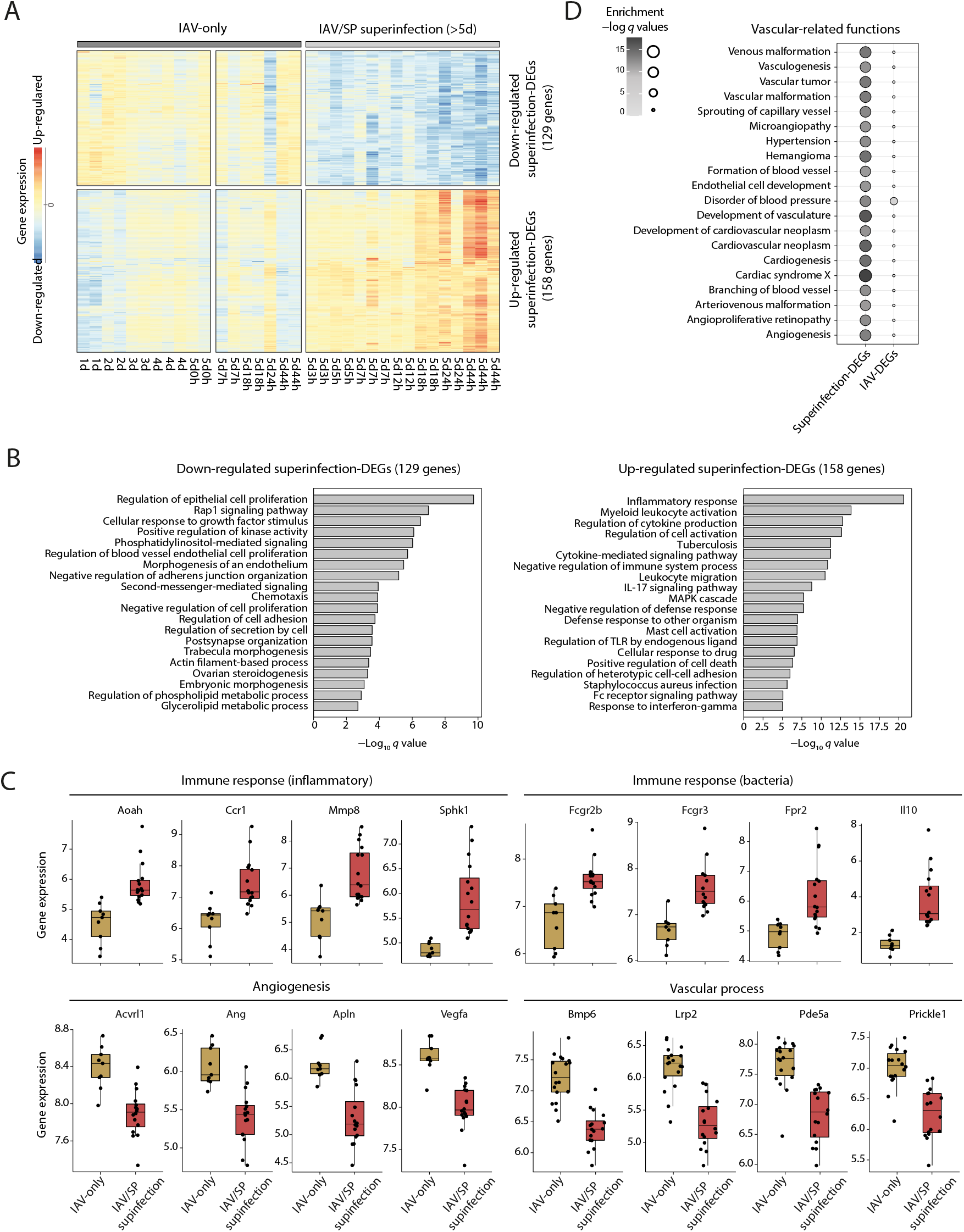
Functional annotation of top superinfection-DEGs. (**A**) Heatmap showing gene expression of top 287 superinfection DEGs. Columns represent a sample (specific treatment and time point) and rows represent genes. The red and blue gradients indicate up- and down-regulated gene expression, respectively, relative to untreated mice. (**B**) Gene ontology (GO) enrichment analysis of the downregulated (left) and upregulated (right) genes from A. Presented are FDR-adjusted -log *q*-values (HG test). (**C**) Boxplots of exemplified genes from A, providing gene expression (log scaled) of vascular, angiogenesis and immune-related genes in the IAV-only and the IAV/SP superinfection groups. Individual samples are shown as dots. (**D**) A bubble plot, demonstrating enrichments in the superinfection-DEGs (left) and IAV-DEGs (right) for various vascular-related functions from the Ingenuity knowledge base annotation. Bubble size and color scale represents the FDR-adjusted HG test *q*-values.

### Reprogramming of the host defense response during superinfection

We next sought to investigate the molecular state of the host resistance program against pathogens. To explore this, we used a predefined transcriptional signature for resistance (Cohn et al., 2022), which allowed us to calculate the ‘resistance level’ in the lungs of each mouse (**Methods**). Using this signature, we found that the general resistance level in response to superinfection is consistent with previous observations (Cohn et al., 2022): (i) resistance was upregulated in response to moderate and severe infections (**Figure 4A**), and (ii) superinfection was marked by higher pathogen and higher resistance levels compared to a single infection (**Figures 4AB, 1DE**), in agreement with the general understanding that the resistance level is coordinated with the pathogen load.

**Figure 4.**
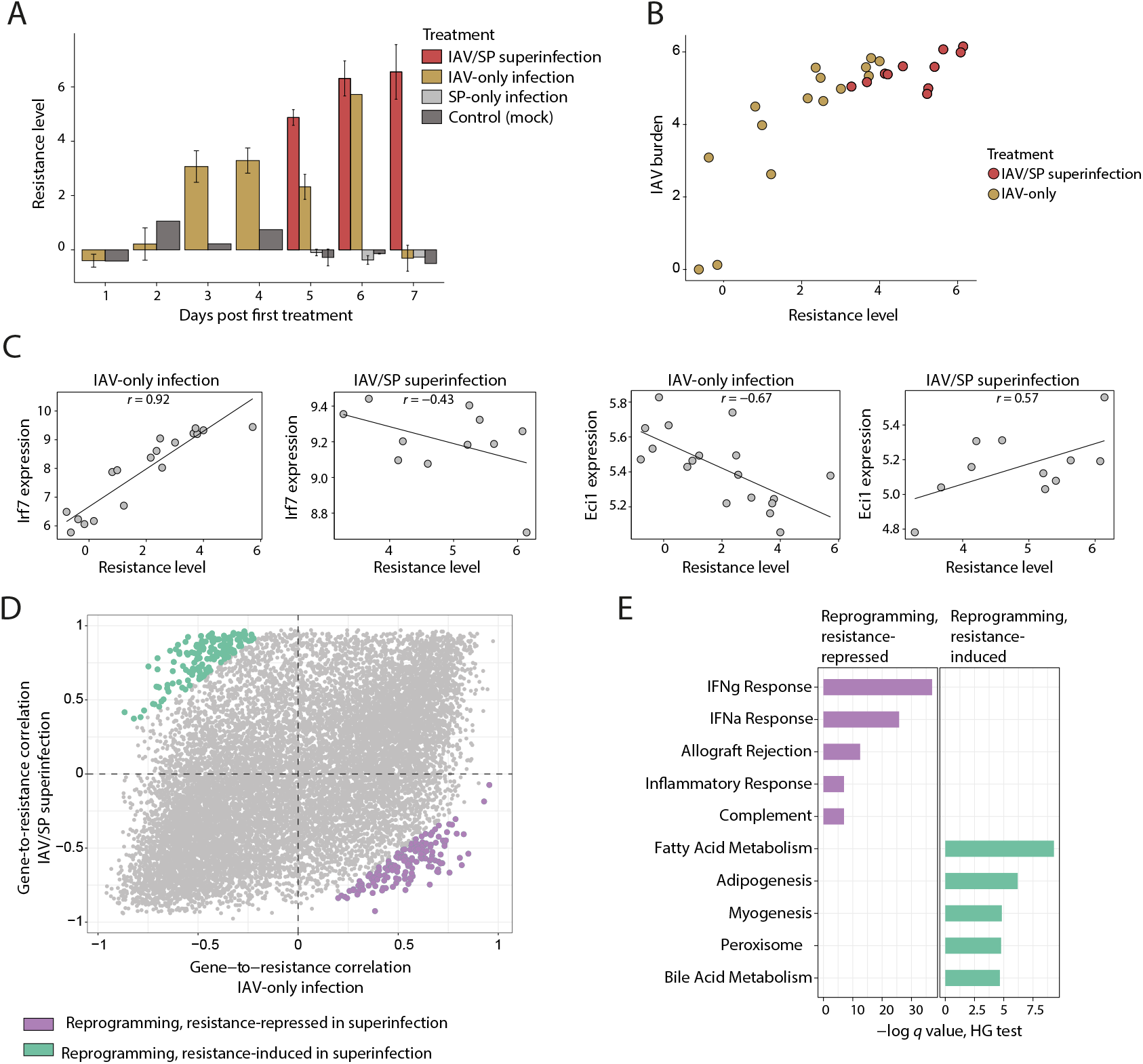
Reprogramming of the anti-pathogen defense (‘resistance’) program in superinfection. (**A**) The host resistance levels (y axis) for individuals in the four treatment groups (color coded) in each time point (x axis). (**B**) A scatter plot of resistance levels (y axis) against the IAV burden (x axis) across individuals (dots). Individuals are color coded according to their treatment group. (**C**) Demonstration of selected genes in which the correlations with resistance are altered between the IAV-only group and the IAV/SP-superinfection group. In each scatter plot, shown are the relation between resistance levels (x axis) and the expression of a gene (*Irf7* or *Eci1*, y axis) across individuals (dots) from a specific treatment group (indicated on top). (**D**) An overview of superinfection-specific changes in gene-to-resistance correlations. For each gene (a dot), the gene-to-resistance correlations are defined across IAV-only mice (x axis) and superinfected mice (y axis). Highlighted are 300 genes with top changes in their resistance-correlations between the IAV-only and superinfected (150 genes in each direction), referred to as ‘reprogrammed genes’ and color coded by their direction of reprogramming. (**E**) Enrichment analysis of the top-reprogrammed genes (resistance-induced and resistance-repressed gene sets are as defined in panel D). Presented are FDR-adjusted HG test *q*-values.

We next analyzed the correlation of each gene with the level of resistance and how this correlation is altered in superinfection. The correlation was calculated across individuals using a separate analysis of the IAV-only and the IAV/SP-superinfection groups. For instance, the antiviral *Irf7* gene has a positive correlation with resistance in IAV-infected mice (Pearson’s *r* = 0.92) but a negative correlation with resistance in superinfection (Pearson’s *r* = – 0.44), suggesting ‘reprogramming’ from positive to negative associations of *Irf7* with the resistance program (**Figure 4C**). We further exemplify *Eci1*, an essential gene for beta-oxidation of unsaturated fatty acids (Hiltunen et al., 2003), with the inverse reprogramming (**Figure 4C**). Thus, *Irf7* is switched from induction to repression by resistance, whereas *Eci1* is switched from repression to induction by resistance. The top 150 genes that are switched from induction to repression by resistance (highlighted in purple in **Figure 4D**) are enriched with interferon signaling genes (HG test *p* <10^-11^, **Figure 4E**; e.g., *Stat2, Irf7, Lgals3bp, Adar, Irf9, Pml*). In contrast, the top 150 genes that are switched from repression to induction by resistance (highlighted in green in **Figure 4D**) are enriched with metabolism-related pathways – for instance, adipogenesis and fatty acid metabolism (HG test *p* < 10^-4^, 10^-2^, respectively, **Figure 4E**; e.g., *Sod1, Cryz, Bckdhb, Gpat4, Acaa2 and Eci1*).

The analysis of superinfection reprogramming has several implications. First, it suggests the reuse of the existing resistance program (for IAV infection) in superinfection, rather than the emergence of a new superinfection-specific program. Second, the analysis suggests metabolic reprogramming as a novel potential process that is involved in superinfection. Third, despite the over-activation of resistance in superinfection, we found that interferon signaling is not overactivated but rather is repressed with increasing resistance levels. We note that both the induction of metabolic reprogramming and the repression of antiviral response have not been previously reported, likely because these effects could only be discerned when considering gene expression relative to the background level of resistance.

### Concluding remarks

Secondary bacterial infections are the most frequent complications during IAV pandemic outbreaks, contributing to excessive morbidity and mortality in the human population (McCullers, 2014). Most IAV-related deaths are attributed to SP infections, which usually begin within the first week of IAV infection in the respiratory tracts (Rynda-Apple et al., 2015; Shrestha et al., 2013).

Many previous studies aimed to explain the enhanced susceptibility to secondary bacterial infections. Early investigations have pointed to lung epithelium damage as a factor in enhanced bacterial adherence and growth ((Brian W. Donnelly et al., 1990; Nugent and Pesanti, 1983; Plotkowski et al., 1986). Other studies have shown that primary IAV infections impair macrophages and neutrophils’ functions in the lung (Ghoneim et al., 2013; McNamee and Harmsen, 2006). Recent reports attributed various changes in cytokines levels, such as TNF-α, IL-6, type I IFN, IL-10, IL-27 (Cao et al., 2014; Gou et al., 2020; McCullers, 2006; Nakamura et al., 2011; van der Sluijs et al., 2004), as well as changes in immune cells composition (e.g., monocytes, T-cells, B-cells (Ellis et al., 2015; Li et al., 2012; Wolf et al., 2014; Wu et al., 2015)), which occur during the viral infection and in turn limit the innate pulmonary host defense against the subsequent bacterial invasion. Although these studies have greatly extended our understanding regarding the interactions of IAV and SP with the host response, systematic identification of potential mechanisms has not been applied.

To determine early events during IAV/SP superinfection, we explored changes in gene expression patterns in superinfected mice compared to IAV-only and SP-only mice. Our data show that superinfected mice exhibited rapid gene expression induction or reduction within the first 12h. Cell proliferation and immune response activation processes were upregulated, while endothelial, vasculogenesis, and angiogenesis processes were downregulated. We detected reduced levels of several angiogenic growth factors, such as VEGF and Apln, which typically induce pro-angiogenic pathways in endothelial cells (Helker et al., 2020; Nowak et al., 2010). These findings support the notion that in addition to the well-characterized impairment of the innate immune response, the downregulated endothelial signature may reflect vascular destruction, which in turn leads to enhanced bacterial dissemination (Sender Vicky et al., 2020).

Our study yields a highly detailed transcriptional profiling that makes it possible to identify subtle changes in the program of the host’s anti-pathogen defense (‘resistance’). Our study suggests that the specific resistance response to superinfection involves reprogramming (from a negative to a positive association with resistance) of fatty acids metabolism. Furthermore, despite the overall hyper-activation of resistance against the pathogen, the specific resistance response to superinfection also involves interferon signaling reprogramming (from a positive to a negative association with resistance). The reprogrammed responses have not been previously reported, likely because these could only be revealed when considering the background state of the host defense. The superinfection-specific reprogramming provides promising targets for future therapeutic interventions.

## Methods

### Mice

Adult female C57BL/6 mice were purchased from Envigo, Israel. Mice were housed under humidity and temperature-controlled specific pathogen-free conditions at the animal facility of Tel Aviv University. Mice were housed on hardwood chip bedding under a 12 h light/dark cycle. Mice were given tap water and standard rodent chow diet ad libitum from their weaning day until the end of the experiment. All animal experiment protocols were approved and conducted following the Institutional Animal Care and Use Committee’s guidelines at Tel Aviv University (approval number 04-17-052).

### Treatment groups

Mouse-adapted H1N1 influenza A/PR/8/34 virus was kindly provided by Dr. Michal Mandelboim (Central Virology Laboratory, Ministry of Health, Chaim Sheba Medical Center) and stored at −80°C. Briefly, the virus was grown in allantoic fluid of 10-day-old embryonated chicken eggs at 37°C for 72 h. Allantoic fluid was harvested, and viral titers were determined by standard plaque assay in Madin–Darby canine kidney (Szretter et al., 2006). SP (ATCC 6303, *S. pneumoniae* type 3 encapsulated strain) was grown for 16 h at 37^0^C in 5% CO_2_ on Columbia Agar plates supplemented with 5% (vol/vol) sheep blood. Colonies were picked and grown in Todd Hewitt broth with yeast extract to mid-logarithmic phase, harvested, and diluted to concentrations of 1 × 10^4^ coloby forming units (CFU) (verified by re-plating 10-fold dilutions). Mice were subjected to four treatments: mock-infection (PBS) control, IAV-only infection, SP-only infection, and IAV infection followed by SP infection (IAV/SP superinfection). Mice were anesthetized intraperitoneally with a ketamine/xylazine cocktail. For the secondary pneumococcal infection model: anesthetized mice were administered intra-nasally with IAV (100 plaque forming units [PFU], tittered on MDCK cells), and five days later, they were infected intra-nasally with SP (1 × 10^4^ CFU). Mice were monitored for additional 44h. Weight and survival were monitored daily. Mice exhibited severe clinical signs of disease or mice with more than 20% weight loss were humanely euthanized.

### Bacterial and viral load in lungs

For bacterial load quantification (by 16S rRNA measurements), total RNA (1 μg) was reverse transcribed using the SuperScript kit (BioRad), and the Real-time PCR was performed in three technical replicates using the Applied Biosystems StepOnePlus Real-Time PCR system and Fast SYBR Green Master Mix (Applied Biosystems). Oligonucleotides for 16S rRNA were:

16s-F: GGTGAGTAACGCGTAGGTAA and 16s-R: ACGATCCGAAAACCTTCTTC.

Relative gene expression levels of the 16S rRNA were normalized relative to the mock-infected samples. IAV load was measured using the viral mRNA levels in the lung tissue based on the bulk transcriptome measurements – particularly, we aligned the RNA-seq reads of all samples against the mouse and the IAV genomes and quantified the viral genes. The reported viral load is the average of viral mRNA across all IAV genes.

### RNA isolation, library preparation, and sequencing

Lung tissues were collected into RNAlater (Qiagen) and lysed with QIAzol (Qiagen). RNA was isolated with miRNeasy kit (Qiagen), and its RNA Integrity Numbers (RINs) were verified to be higher than 8 using the Agilent 2100 Bioanalyzer. cDNA libraries were prepared using 2μg of the isolated RNA and the SENSE mRNA-Seq Library Prep Kit V2 for Illumina (Lexogen). DNA size and quality were checked using the Agilent 2100 Bioanalyzer. Libraries were quantified using the Qubit DNA HS Assay kit (Invitrogen). The amplified libraries, each sample having its unique index primer, were pooled at a total concentration of 2 nM and sequenced using the Illumina HiSeq platform (Technion Genome Center, Israel). Clustering, demultiplexing, and alignment were performed as previously described (Ushakov et al., 2017). This dataset is deposited in the GEO database (GSE206534).

### RNA-Seq data analysis

Raw counts were derived with featureCounts (Liao et al., 2014). Raw read counts were imported into R studio (version 3.6.1) and normalized using the DESeq2 package (Love et al., 2014, p. 2). Reported DEGs for each comparison obtained *q* values (i.e., FDR-adjusted p-values) that are lower than 0.05 and FC levels that are higher than 1.5 (either for up- or down-regulation) (**Supp. Table 1**). To select the top superinfection-DEGs, we first clustered all superinfection-DEGs and then selected the genes included in the two clusters with the strongest superinfection-specific response (clusters are shown in **Figure 3A** and detailed in **Supp. Table 1**). Enrichment analyses were performed using functional classes in the Gene Ontology (GO), Ingenuity Pathway analysis (IPA) and Enrichr (Kuleshov et al., 2016) (multiple testing FDR correction was applied in all cases). Resistance levels were calculated by a predefined linear combination of all genes (Cohn et al., 2022). Before calculating the resistance levels, the expression levels of each gene were centered according to the measurements in the mock-infected mice.

## Supplementary Tables

**Supplementary Table 1. Differentially expressed genes.** In columns 1-13, for each of the comparisons between treatment groups (row 1), indicated are the differential expression *q* values, log2 FC levels, and indications of DEGs (row 2). Superinfection-DEGs and top superinfection-DEGs are indivated in columns 14 and 15, respectively.

## Acknowledgments

This work was supported by by European Union Horizon 2020 under grant agreement No. 847422 (ImmunoSep). IG-V is a Faculty Fellow of the Edmond J Safra Center for Bioinformatics at Tel Aviv University. GY was supported by the Edmond J Safra Center for Bioinformatics at Tel Aviv University.

## Notes

### Competing Interest Statement

The authors have declared no competing interest.

